# Acute Vigorous Exercise Decreases Subsequent Non-Exercise Physical Activity and Body Temperature Linked to Weight Gain

**DOI:** 10.1101/2023.10.25.563892

**Authors:** Daisuke Funabashi, Shohei Dobashi, Kazuki Sameshima, Hiroyuki Sagayama, Takeshi Nishijima, Takashi Matsui

## Abstract

**Purpose:** Exercise benefits the body and mind, but its weight loss effect is less than generally expected. Although this phenomenon is likely primarily due to a decrease in non-exercise physical activity (NEPA) resulting in a decrease in non-exercise activity thermogenesis, the underlying mechanisms and effects of exercise intensity remain unknown. Here we show that acute vigorous exercise decreases subsequent NEPA and body temperature (BT) in association with body weight gain.

**Methods:** Adult male C57BL/6J mice were categorized into three groups: sedentary, moderate exercise, and vigorous exercise, with exercise groups undergoing a 30 min treadmill session. Using an intraperitoneally implanted activity monitor, NEPA and BT were monitored for two days before and three days after exercise. The daily synchrony between NEPA and BT was evaluated using a cross-correlation function. Plasma corticosterone was also detected 6 and 24 h after exercise.

**Results:** Notably, Only the vigorous exercise group exhibited a decline in both NEPA and BT, resulting in body weight gain the following day, despite no observed changes in food intake. Furthermore, vigorous exercise induces a distinct delay in the daily dynamics of NEPA compared to BT. A positive correlation was observed between plasma corticosterone levels and changes in NEPA levels before and after exercise across all exercise groups.

**Conclusions:** Our findings provide evidence for vigorous exercise-specific reduction in subsequent NEPA, BT, and their synchrony linked to weight gain, likely due to the disturbed circadian rhythm of corticosterone. This ultimately redefines the significance of exercise intensity in beneficial effects beyond the energy expenditure of the exercise itself.

## Introduction

Exercise has been widely recommended as a pivotal strategy to improve physical and mental health (1–3). Given the growing prevalence of obesity, a leading cause of many health problems, optimal exercise prescriptions are required to achieve better energy balance and prevent obesity. However, the effect of exercise on body weight loss is often less than the theoretically predicted reduction based on exercise-induced energy expenditure (4,5). This is considered to be because exercise-induced increases in energy expenditure reduce the energy spent on other physiological activities (6). Therefore, we should better understand not only the effects of exercise itself, but also the physiological and behavioral post-effects of exercise to develop a comprehensive exercise strategy for preventing obesity and promoting a healthy lifestyle.

Non-exercise components related to energy expenditure include non-exercise physical activity (NEPA) and non-exercise activity thermogenesis (NEAT) in association with NEPA. NEPA refers to almost all activities in daily life, namely unconscious and nonvolitional movements, excluding structured and purposeful exercises. Energy expenditure resulting from NEPA, known as NEAT, is the most variable component of total energy expenditure (7). Consequently, NEPA and the resulting NEAT are closely associated with fat gain and obesity risk (1). In addition, a reduction in NEPA may predispose individuals to physical inactivity even if they engage in exercise. Notably, low levels of NEPA are implicated in the deterioration of cognitive function and increased risk of dementia (8–10). In light of these findings, maintaining optimal levels of NEPA and the resulting NEAT is required for fostering overall health, including preventing obesity and maintaining mental health.

Exercise induces a compensatory reduction in NEPA following exercise itself, and this phenomenon is a critical contributor to a lack of weight loss after exercise than expected theoretically (11–13). In rodents, the introduction of a running wheel decreases NEPA (14–16), underscoring the potential negative impact of even voluntary exercise on NEPA. Additionally, a single bout of treadmill running decreases NEPA levels following exercise in mice (17). These findings collectively indicate that exercise decreases subsequent NEPA. However, the effects of exercise type, intensity, and volume on post-exercise compensatory reduction in NEPA and NEAT and thermogenesis resulting from other physiological activities remain largely unclear.

The recommendation of moderate to vigorous physical activity for overall health is well established, while the benefits of exercise vary depending on exercise intensity (18,19). Despite the acknowledged efficacy of vigorous exercise in enhancing metabolic health (20), it sometimes fails to improve spatial learning and anxiety-like behaviors and increases the expression of mature brain-derived neurotrophic factor proteins in the hippocampus (21,22). This may stem from the excessive stress-related physiological responses induced by vigorous exercise, such as the activation of corticotrophin-releasing factor neurons in the hypothalamic paraventricular nucleus and increases in corticosterone release in rodents (23,24). Furthermore, corticosterone exhibits circadian rhythms that peak at the beginning of the active phase, a dark period in rodents, and synchronize with the dynamics of physical activity (25,26). Furthermore, plasma corticosterone levels were positively correlated with wheel-running distance in mice (27), suggesting an interplay between corticosterone and physical activity levels. Given the specific increase in corticosterone following vigorous exercise and the close relationship between physical activity and corticosterone, post-exercise compensatory reduction in NEPA and NEAT may also be triggered by vigorous exercise in association with corticosterone.

However, human studies face difficulties in accurately and continuously detecting NEPA and core body temperature as indicators of whole-body thermogenesis after exercise (28,29). Indeed, compensatory responses in NEPA to exercise intervention often exhibit considerable variability among participants due to challenges in controlling external factors such as genetic and social background, living conditions, dietary habits, and other free-living activities (30,31). In addition, ingestible telemetric capsules are often used for the measurement of core body temperature in free-living humans (32), but accurate detection could fail owing to the influence of some foods/drinks (29). It is also difficult to measure for a long duration over several days, since ingested capsules are excreted in about 30 hours (29,33). Consequently, these methodological difficulties contribute to a poor understanding of compensatory physiological responses following exercise.

We thus employed an animal model realizing for accurate- and continuous-measurement of NEPA and core body temperature simultaneously (34,35), and tested the hypothesis that acute vigorous, but not other intensity, exercise decreases subsequent NEPA and core body temperature. We first performed simultaneous measurements of NEPA, core body temperature, and corticosterone levels following different exercise intensities in mice. To examine the effects on physiological circadian rhythms, we next analyzed the interindividual synchrony between NEPA and core body temperature following exercise. Finally, to examine the mechanism of the compensatory reduction in NEPA following exercise, we analyzed the relationship between NEPA, body temperature, body weight, and corticosterone levels.

## Methods Animals

Male C57BL/6 mice (11 weeks old; n = 27) (SLC, Shizuoka, Japan) were used in these experiments. Three mice were excluded due to an accident during the experiment and the resulting change in group size. The mice were housed in standard laboratory cages (225 × 338 × 140 mm) under controlled temperature (22 ± 2 °C) and light (12:12-h light and dark cycle, light at 7:00– 19:00). Food and water were provided ad libitum. All experimental protocols were conducted in accordance with the guidelines of the University of Tsukuba Animal Experiment Committee.

### Experimental designs

A schematic of the experimental procedure is shown in Figure 1A. After 1-week habituation to the housing environment, mice underwent surgery to implant a device intraperitoneally to measure NEPA and body temperature, as described previously (34). After the 1-week of recovery, all mice were habituated to running on a treadmill (Natsume, Tokyo, Japan) for five sessions over 6 days for 30 min/day with incremental increases in speed (from 0 to 25 m/min). After the habituation period, all mice were randomly divided into three groups: sedentary (SE), moderate (ME), and vigorous (VE) exercise, and exercised on a treadmill for 30 min at each intensity. Based on a previous study that determined the lactate threshold (LT) during treadmill running in mice (36), treadmill running was performed at SE (0 m/min), ME (LT, 15 m/min), and VE (supra-LT, 25 m/min). The NEPA and core body temperature were continuously measured in the home cage (Fig. 1B) from 2 days before to 3 days after the exercise intervention, using an intraperitoneally implanted device. Three days after the first exercise intervention, the mice performed the same treadmill running again, and blood was sampled 6 and 24 h after exercise to measure plasma corticosterone levels.

**Figure 1.**
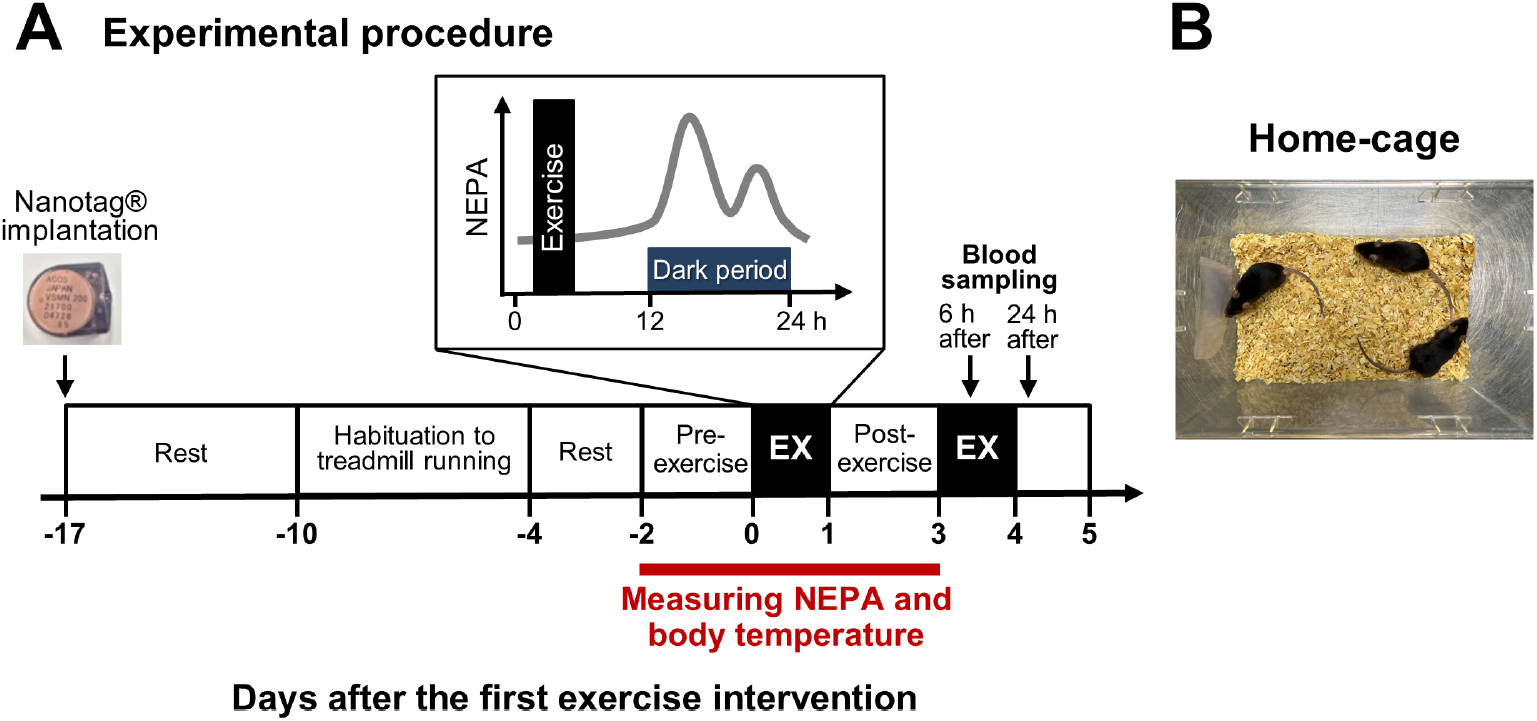
Schematic overview of this experiment. Schematic of the experimental procedure (A). Exercise was performed on the day of the exercise as an EX. Images of the home cage in which mice were housed throughout this experiment including during the measurement of NEPA and body temperature (B).

### Measuring NEPA and body temperature

NEPA and core body temperature were measured using nanotag® (Kissei Comtec Ltd, Japan), a recently developed body-implantable device (34). Mice aged ten weeks were implanted with nanotag® intraperitoneally under anesthesia via the intraperitoneal injection of an anesthetic cocktail (hydrochloric acid medetomidine, 0.3 mg/kg; midazolam, 4 mg/kg; butorphanol tartrate, 5 mg/kg). After surgery, the mice were intraperitoneally administered atipamezole hydrochloride (an antagonist of medetomidine hydrochloride, 0.3 mg/kg) to recover from anesthesia, thereby preventing a sustained decline in body temperature. The mice underwent a 1-week recovery period after surgery because the negative effect on physical activity was nearly negligible after this period (34). Prior to nanotag® implantation, we set the device to start and stop recording automatically according to the experimental procedure (Fig. 1A). The number of counts was recorded in the nanotag ® at 5-minute intervals. After the experiment was completed, the nanotag ® was removed from the deeply anesthetized mice, and the data stored in the nanotag® were retrieved into a computer using the FeliCa® communication system.

### Synchrony analysis

Nanotag® allows us to continuously measure NEPA and body temperature simultaneously in mice. These data were used to calculate the cross-correlation function (CCF) between daily changes in NEPA and body temperature on day 1 post-exercise, which can detect the strength of correlations and time lags among the temporal data. The time lag representing the peak CCF was obtained for each participant to investigate the causality between daily changes in NEPA and body temperature. A negative time lag indicated that the circadian rhythm of body temperature preceded that of the NEPA.

### Corticosterone

Corticosterone concentrations in mouse plasma samples were measured using the Corticosterone ELISA Kit (ARB, #K014-H1) according to the manufacturer’s recommended protocols. Absorbance was determined using a microplate reader (Varioskan LUX Multimode Microplate Reader, Thermo Fisher Scientific, MA, USA).

### Statistical analysis

Daily changes in NEPA and body temperature were analyzed using a two-way (time × exercise-intervention) repeated-measures analysis of variance (ANOVA). Percent changes in NEPA and body temperature from pre-exercise to days 1–3 post-exercise and body weight before and 24 hours after exercise were analyzed using two-way (time × group) repeated-measures ANOVA. If a significant interaction or main effect was observed, Tukey’s or Dunnett’s multiple comparison test was performed. The CCF analysis (window size of 30 data points) was performed using the original program in Python 11. Percent changes in NEPA and body temperature during the light and dark periods from pre-exercise to day 1 post-exercise, body weight changes, plasma corticosterone levels, CCF, and lag were analyzed using ordinary one-way ANOVA. Pearson’s correlation analysis (percentage changes in NEPA vs. percentage changes in body temperature, percentage changes in NEPA vs. body weight change, body temperature vs. body weight change, and plasma corticosterone levels vs. percentage changes in physical activity) was also performed. All data are presented as the mean ± standard error (SEM). The threshold for statistical significance was set at P < 0.05.

## Results

### Vigorous exercise decreases subsequent non-exercise physical activity

We first compared the daily changes in NEPA during pre-exercise with that on day 1 post-exercise within each group (Fig. 2A–C). Although there was no time × exercise-intervention interaction [F (19, 95) = 0.925, P = 0.554] in the SE group (Fig. 2A), the ME and VE groups showed a time × exercise-intervention interaction [F (19, 152) = 1.779, P < 0.05; F (19, 152) = 4.470, P < 0.0001]. In ME, we observed a significant difference in NEPA at only one time point between pre-exercise and day 1 post-exercise, whereas VE exhibited significant differences at some time points (Fig. 2B, C).

**Figure 2.**
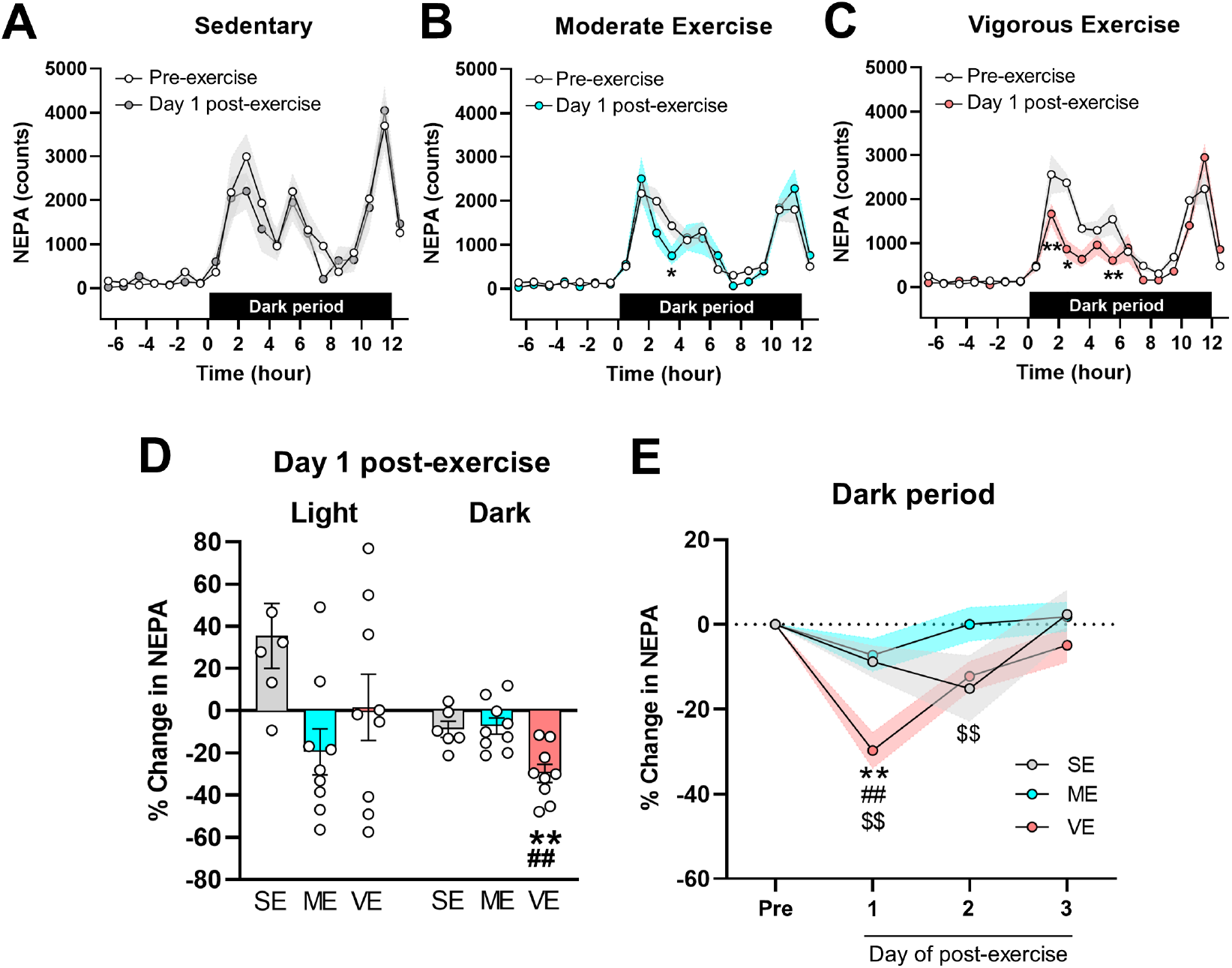
Vigorous exercise decreases subsequent non-exercise physical activity. Daily changes in NEPA pre-exercise and on day 1 post-exercise in SE (A), ME (B), and VE (C). The pre-exercise NEPA was averaged for two days of NEPA before exercise. ^*^P < 0.05, ^**^P < 0.01 vs. pre-exercise. Percentage change in NEPA from pre-exercise to day 1 post-exercise during the light and dark periods (D) ^**^P < 0.01 vs. SE. Percentage change in NEPA during the dark period from pre-exercise to days 1–3 post-exercise (E). ^**^P < 0.01 vs. SE, ^# #^P < 0.01 vs. ME, ^$$^P < 0.01 vs. pre-exercise.

The percentage change in NEPA from pre-exercise to day 1 post-exercise was calculated during the light and dark periods, respectively, and compared among groups (Fig. 2D). A significant main effect of group was observed in the percentage change in NEPA during the dark period (P < 0.01), but not in the light period. During the dark period, the percentage change in NEPA in the VE group was significantly lower than that in the SE and ME groups (P < 0.01). Next, in the analysis of the percentage change in NEPA during the dark period from pre-exercise to days 1–3 post-exercise (Fig. 2E), two-way repeated-measures ANOVA revealed a significant time × group interaction [F (6, 63) = 4.87, P < 0.001]. In the post hoc analysis, a significant reduction from pre-exercise to days 1 and 2 post-exercise was observed only in the VE group (P < 0.01). These results clearly indicate that post-exercise NEPA levels decrease in a vigorous exercise-specific manner.

### Vigorous exercise leads to a subsequent reduction in body temperature during the active dark period

Daily changes in body temperature were compared between pre-exercise and day 1 post-exercise in each group (Fig. 3A–C). In the SE group, daily changes in body temperature did not differ between pre- and post-exercise [F (19, 95) = 1.39, P = 0.152]. In contrast, the ME and VE showed a time × exercise-intervention interaction [F (19, 152) = 4.14, P < 0.0001; F (19, 152) = 5.34, P < 0.0001]. Post hoc analysis revealed a significant decline in body temperature on day 1 post-exercise compared with pre-exercise at some time points during the dark period in the ME and VE groups (Fig. 3B, C).

**Figure 3.**
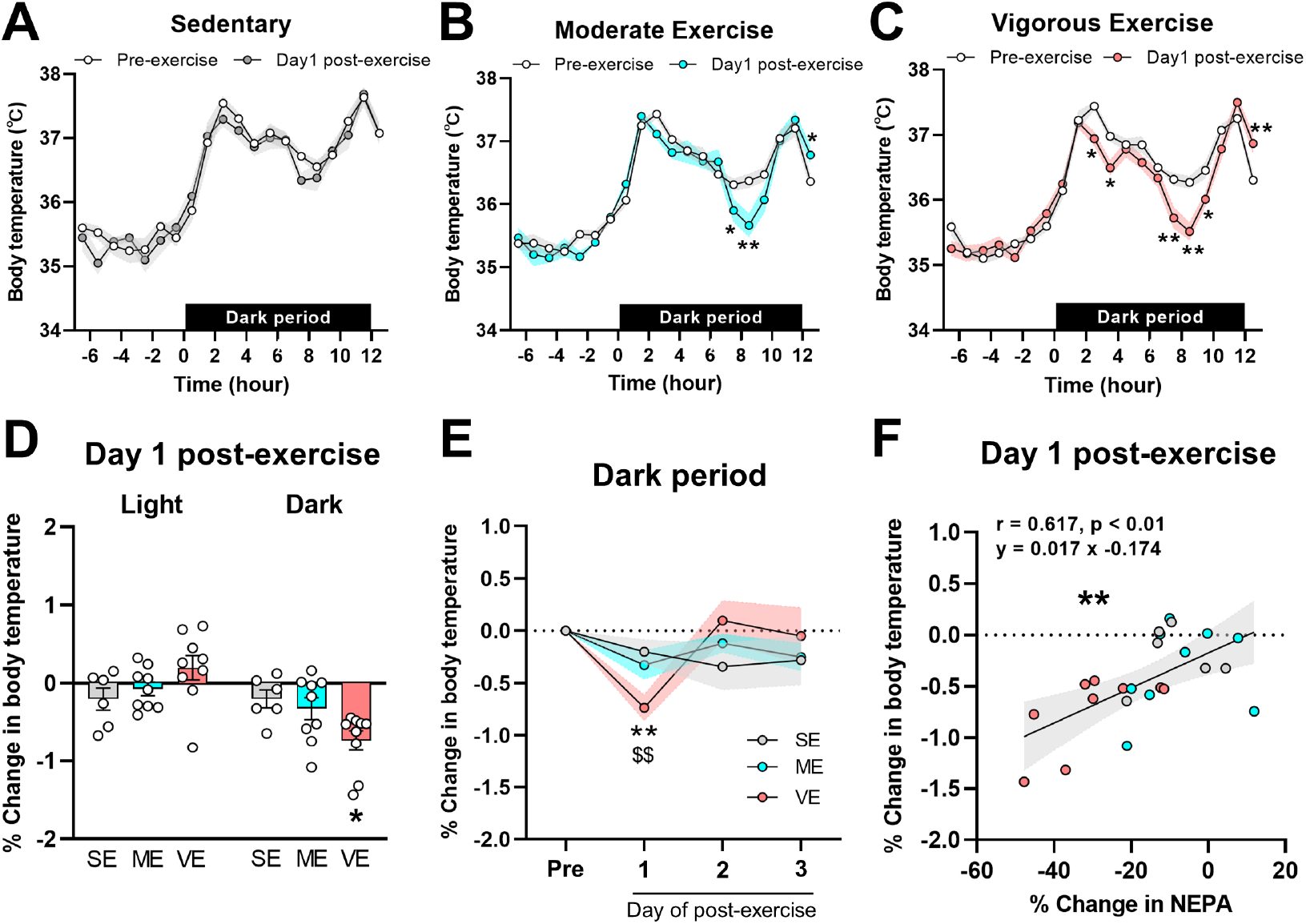
Vigorous exercise leads to a subsequent reduction in body temperature during the active dark period, related to the change in non-exercise physical activity. Daily changes in body temperature pre-exercise and on day 1 post-exercise in SE (A), ME (B), and VE (C). Pre-exercise body temperature was averaged for two days of body temperature before exercise. ^*^P < 0.05, ^**^P < 0.01 vs. The percent change in body temperature from pre-exercise to day 1 post-exercise during the light and dark period (D) ^*^P < 0.05 vs. SE. Percentage change in body temperature during the dark period from pre-exercise to days 1–3 post-exercise (E). ^**^P < 0.01 vs. SE, ^$$^P < 0.01 vs. pre-exercise. Correlation between the percentage change in NEPA and body temperature from pre-exercise to day 1 post-exercise (F). The 95% confidence bands are shown in gray-shaded areas. ^**^P < 0.01.

The percentage change in body temperature from pre-exercise to day 1 post-exercise was calculated during the light and dark periods, respectively, and compared among groups (Fig. 3D). One-way ANOVA revealed a significant main effect of group during the dark period (P < 0.05), but not during the light period. During the dark period, the percentage change in body temperature from pre-exercise to day 1 post-exercise in the VE group was significantly lower than that in the SE group (P < 0.05). In the analysis of the percentage change in body temperature during the dark period from pre-exercise to days 1–3 post-exercise (Fig. 3E), two-way repeated-measures ANOVA revealed a significant time × group interaction [F (6, 63) = 2.89, P < 0.05]. We observed a post-exercise reduction in body temperature in the VE group (P < 0.01), and the percent reduction at day 1 post-exercise was larger in the VE group than in the SE group (P < 0.05). In addition, we observed a significant positive correlation between the percentage change in NEPA and body temperature from pre-exercise to day 1 post-exercise (r = 0.617, P < 0.01, Fig. 3F). These results provide evidence for a vigorous exercise-specific decrease in post-exercise body temperature associated with NEPA levels, implicating disrupted thermogenesis.

### Vigorous exercise increases body weight without altering food intake

Body weights were compared before and 24 h after exercise in each group. Since two-way repeated-measures ANOVA revealed a significant time × group interaction [F (2, 20) = 7.586, P < 0.01] (Fig. 4A), we additionally compared body weight changes among groups (Fig. 4B). Body weight change was significantly greater in the VE group than in the SE group (P < 0.01), suggesting that vigorous exercise facilitated subsequent body weight gain. Food intake for 24 hours following exercise was calculated per cage and divided by the number of mice, because mice were housed in a group (three mice per cage) in our experiment. Average 24-hour food intake was comparable among groups: 2.42 ± 0.09 g (SE), 2.49 ± 0.13 g (ME), 2.55 ± 0.08 g (VE).

**Figure 4.**
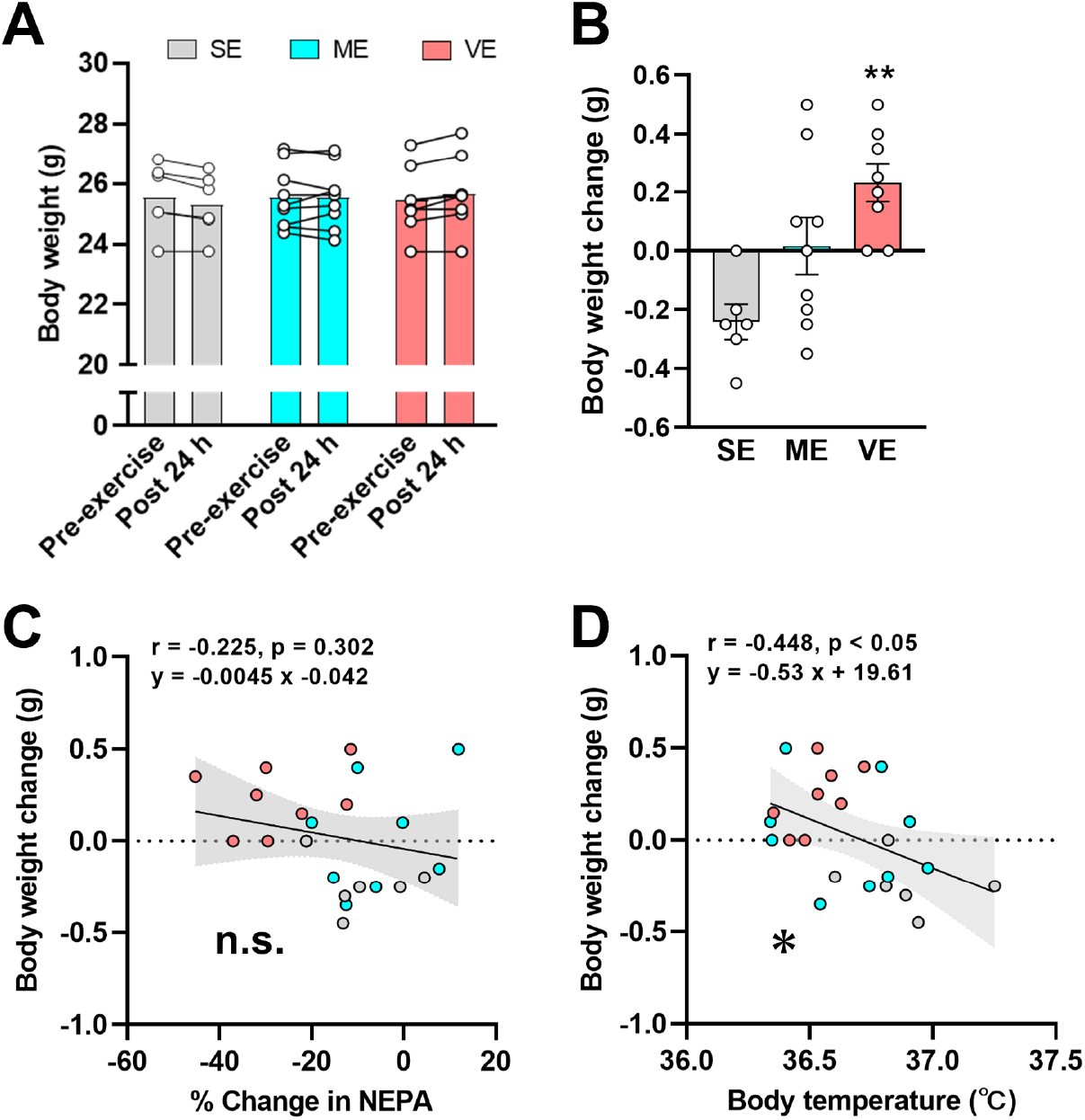
Vigorous exercise increases body weight in association with a decline in body temperature. Body weight before and 24 hours after exercise (A). Body weight changes 24 hours post-exercise (B). ^**^P < 0.01 vs. SE. The correlation between percent change in NEPA from pre-exercise to day 1 post-exercise and body weight changes 24 hours post-exercise (C). Correlation between body temperature on day 1 post-exercise and body weight changes 24 h post-exercise (D). The 95% confidence bands are shown in the gray-shaded areas. ^*^P < 0.05.

Additionally, we investigated whether NEPA and body temperature were associated with body weight changes following exercise. Although there was no correlation between the percent change in NEPA and body weight change (r = -0.225, P = 0.302, Fig. 4C), the average body temperature during the dark period on day 1 post-exercise was significantly correlated with body weight change (r = -0.448, P < 0.05, Fig. 4D). These results suggest the possibility that vigorous exercise reduces body temperature, likely due to lowered thermogenesis, leading to body weight gain the next day without food intake changes.

### Vigorous exercise induces a distinct negative time lag between the daily dynamics of non-exercise physical activity and body temperature

Typical data showed daily changes in NEPA and body temperature during day 1 post-exercise in the ME and VE groups, respectively (CCF = 0.683, Lag = 10 min, Fig. 5A; CCF = 0.641, Lag = -10 min, Fig. 5B). There was no significant difference in the CCF values among the groups (P = 0.391, Fig. 5C). In contrast, the lag time in VE was significantly shifted in the negative direction compared to SE and ME (P < 0.01, P < 0.05, Fig. 5D). This analysis reveal that vigorous exercise induced a distinct delay in the daily dynamics of NEPA compared with body temperature.

**Figure 5.**
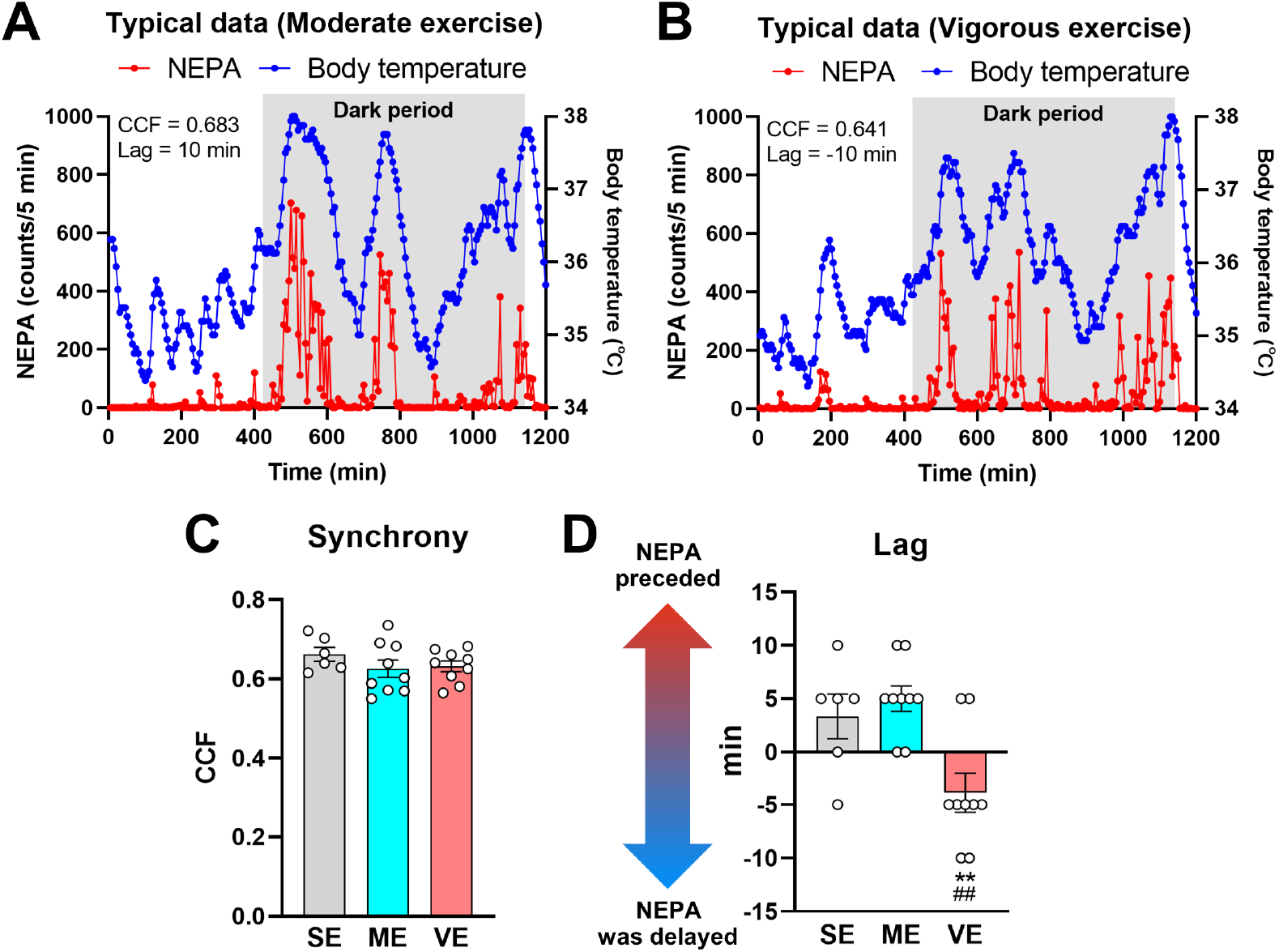
Vigorous exercise induces a distinct negative time lag between the daily dynamics of non-exercise physical activity and body temperature. Typical data of daily changes in NEPA and body temperature during day 1 post-exercise in the ME (A) and VE (B) groups. CCF (C) and lag time (D) of daily changes in NEPA and body temperature on day 1 post-exercise. A negative time lag indicates that the circadian rhythm of NEPA is delayed from that of body temperature. ^**^P < 0.01 vs. SE, ^##^P < 0.01 vs. ME.

### Plasma corticosterone levels are positively correlated with changes in NEPA following exercise in all the exercise groups

No difference was found in plasma corticosterone levels 6 hours after exercise among the groups (P = 0.160, Fig. 6A). There were also no differences in plasma corticosterone levels 24 hours after exercise among the groups (P = 0.138). On the other hand, plasma corticosterone levels 6 hours after exercise were significantly correlated with the percent changes in NEPA from pre-exercise to day 1 post-exercise in the ME (r = 0.819, P < 0.05) and VE (r = 0.897, P < 0.05), respectively (Fig. 6B, C).

**Figure 6.**
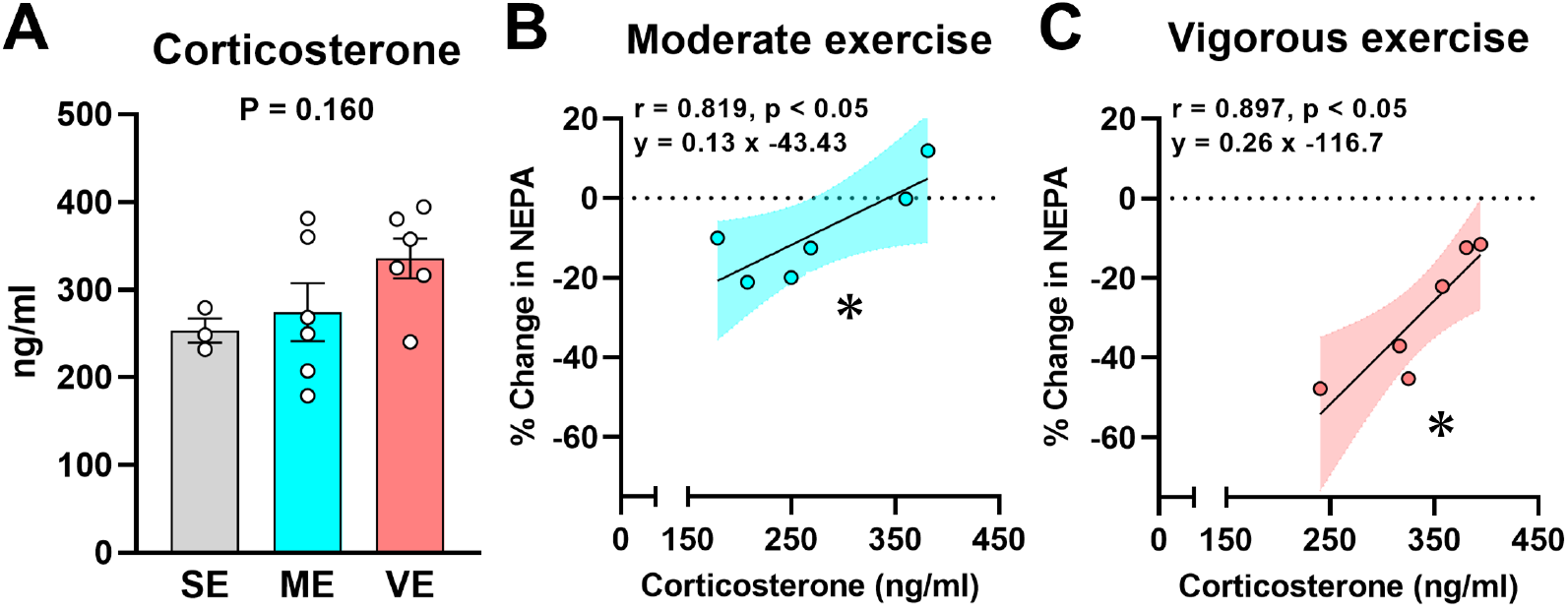
Plasma corticosterone levels were positively correlated with changes in non-exercise physical activity following exercise in all exercise groups. Plasma corticosterone levels 6 hours after exercise (A). Correlation between plasma corticosterone 6 h after exercise and percent changes in NEPA from pre-exercise to day 1 post-exercise in ME (B) and VE (C) groups. The 95% confidence bands are shown for ME (light blue) and VE (pink). ^*^P < 0.05.

## Discussion

In the present study, we examined whether a reduction in NEPA and body temperature and disturbed their synchrony following exercise is specifically caused by vigorous exercise, in association with corticosterone before awakening. In addition, we explored the association between body weight gain and compensatory reduction in NEPA and body temperature following exercise. Our comprehensive findings, summarized in Figure 7, illuminate that vigorous exercise distinctly triggers a compensatory response, characterized by a reduction in NEPA, body temperature, and disturbance of their synchrony. These compensatory responses likely underlie the observed vigorous exercise-induced body weight gain. Furthermore, plasma corticosterone levels prior to awakening are positively correlated with changes in NEPA following exercise, suggesting the potential role of insufficient pre-awakening corticosterone elevation in contributing to compensatory NEPA reduction post-exercise. Our study sheds light on the specific effects of vigorous exercise on NEPA, body temperature, and their synchrony, providing insights into the compensatory responses and subsequent body weight changes following exercise.

**Figure 7.**
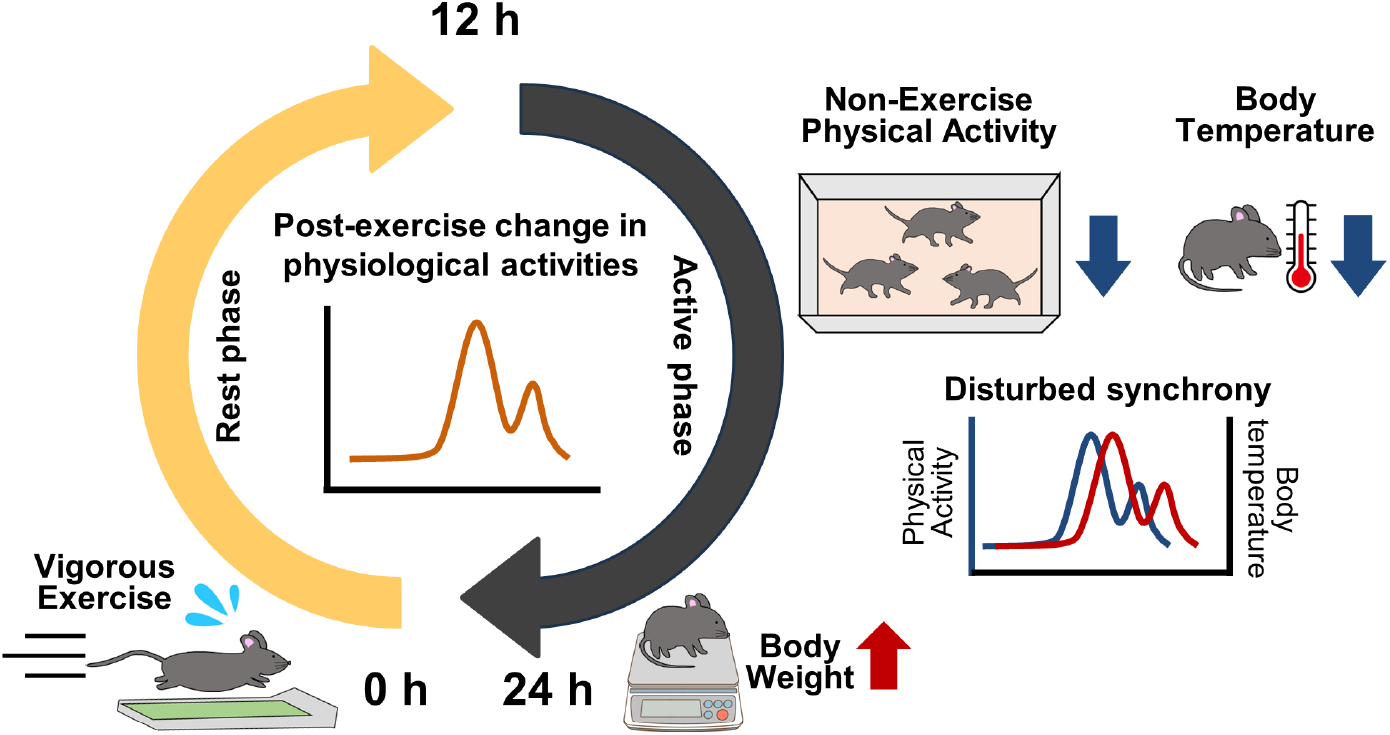
Summary image of the vigorous exercise-specific compensatory physiological responses linking to body weight gain in mice. Vigorous exercise decreases NEPA and body temperature during the active phase and disturbs their synchrony, contributing to body weight gain 24 hours post-exercise in mice.

### Why is weight loss effect of exercise less than generally expected?: Contributions of NEPA and body temperature

So far, growing evidence suggests that the effect of exercise on body weight loss is less than expected (4,5). Pontzer et al. demonstrated that increasing physical activity does not simply add total energy expenditure in a dose-response manner (37). In addition, an animal study reported that the total energy expenditure does not increase despite the fact that the daily running distance increases considerably in mice (16). These findings suggest the possibility of exercise-induced compensatory responses related to the regulation of the energy balance. While prior studies have reported a compensatory reduction in NEPA after exercise (14,15), an exploration into whether the volume or intensity of exercise contributes to the observed decrease in NEPA has remained unexplored. The present study, for the first time, elucidated that NEPA decreases following vigorous exercise but not moderate exercise (Fig. 2E). This implies that a compensatory reduction in NEPA following exercise is specifically caused by vigorous exercise. These novel findings suggest that vigorous exercise may induce a post-exercise state of physical inactivity in mice, emphasizing the potential benefits of moderate exercise as an optimal program for fostering a physically active lifestyle and preventing obesity.

Body temperature, serving as a sensitive indicator of whole-body metabolic activity, immune function, and physical activity levels, is largely influenced by NEAT and basal metabolism (38,39). Therefore, a compensatory reduction in these physiological activities following exercise is more likely to be directly linked to a reduction in body temperature. In this context, we highlighted body temperature as a marker of thermogenesis encompassing NEAT and basal metabolism. Our study revealed the vigorous exercise-specific reduction in body temperature during the dark period following exercise (Fig. 3E). This observation suggests that vigorous exercise leads to a decrease in subsequent thermogenesis, encompassing NEAT and basal metabolism. Intriguingly, we also observed a post-exercise reduction in body temperature during the late dark period (Fig. 3B, C), a time when exercise-induced reduction in NEPA did not occur (Fig. 2B, C). This implies that the post-exercise reduction in thermogenesis during the late dark period is attributed to decreased basal metabolic rate but not to changes in NEAT. Considering that this reduction in body temperature during the late dark period is observed consistently in both ME and VE groups, our results suggest that exercise is likely to elicit a compensatory reduction in basal metabolic rate in mice, independent of exercise intensity. This insight hints at the notion that a decrease in basal metabolic rate may preferentially occur as a compensatory response following exercise compared to a reduction in NEPA.

In our investigation, a notable disparity in body weight gain emerged, with the VE group exhibiting significantly higher weight gain compared to the SE, despite the likelihood of the highest energy expenditure during exercise within the VE group (Fig. 4B). Additionally, post-exercise body temperature negatively correlated with body weight changes (Fig. 4D), indicating that a lower body temperature following exercise leads to body weight gain in mice. This finding emphasizes the role of post-exercise thermoregulation in body weight management. Moreover, our findings highlight a connection between post-exercise reduction in NEPA and a decline in body temperature (Fig. 3F). This suggests that the observed reduction in NEPA may also play a role in contributing to body weight gain in mice. These insights into the potential role of compensatory reduction in NEPA and thermogenesis in post-exercise transient weight gain would help us to develop an optimal program for fostering a physically active lifestyle and preventing obesity.

We observed no body weight reduction or increased food intake in both exercising groups, emphasizing that a reduction in NEPA and body temperature plays a key role in energy compensation following exercise. On the other hand, body weight gain was not different between the SE and ME groups (Fig. 4B), despite the fact that basal metabolic rate was assumed to decline the ME following exercise. This finding may suggest that a reduction in NEPA largely contributes to energy compensation and body weight gain compared with a decline in basal metabolic rate.

Altogether, our findings may propose that exercise initially triggers a compensatory reduction in basal metabolic rate, followed by a subsequent compensatory reduction in NEPA in response to heightened exercise intensity. This insight underscores the significance of exercise intensity in understanding the behavioral and physiological adaptations to exercise. Nevertheless, it is crucial to acknowledge that our study lacked an assessment of energy expenditure. Hence, further investigations are warranted to examine the effects of each exercise intensity on post-exercise metabolic rate assessed by respiratory analysis are required.

### Disturbed circadian rhythm following vigorous exercise: Evidence from interindividual physiological synchrony

NEPA and body temperature exhibit circadian rhythms that are highly synchronized; for example, the first increase in these levels is observed during the early dark period (40). Since these circadian rhythms are likely to interact with each other (38), we attempted to analyze the synchrony in circadian rhythms between NEPA and body temperature. Investigating whether their synchrony is altered following exercise promotes an integrative understanding of the effects of exercise on circadian rhythms. As a result of this synchrony analysis, the CCF between daily changes in NEPA and body temperature on day 1 post-exercise did not differ among groups (Fig. 5C), whereas the time lag significantly shifted toward a negative direction in the VE compared with SE and ME (Fig. 5D). The time lag in the SE and ME indicates that the circadian rhythms of NEPA precede those of body temperature, meaning that an increase in NEPA is generally followed by an elevation of body temperature. In contrast, our results indicated that vigorous exercise delayed the circadian rhythm of NEPA from that of body temperature. This reversal of circadian rhythms between NEPA and body temperature may represent a disturbed biological rhythm in mice.

The function of circadian rhythms is assessed by the amplitude of the rhythm and the timing of the acrophase of that rhythm. For example, aging decreases the amplitude of the rhythm of voluntary physical activity and is linked to the phase advancement of the rhythms of body temperature, cortisol, and melatonin (41–43). To date, a lot of studies have examined the effects of several biological and environmental factors, including aging, exercise, and diet, on circadian rhythms of physical activity and body temperature (44–46). However, the synchrony between the circadian rhythm of physical activity and body temperature has been poorly investigated, and the implications of this synchrony on health-related outcomes remain unknown. In the present study, we observed for the first time that synchrony between the circadian rhythms of NEPA and body temperature is disturbed by vigorous exercise (Fig. 5D). Considering the vigorous exercise-specific reduction in NEPA and body temperature following exercise observed in this study, the synchrony of these circadian rhythms is likely to play a key role in maintaining NEPA and body temperature at optimal levels. For a detailed investigation in the next step, NEPA and body temperature should be measured simultaneously at shorter intervals than those measured in the present study.

### Possible underlying mechanisms of compensatory reduction in NEPA following exercise

We first focused on corticosterone, a hormone that represents a circadian rhythm similar to that of physical activity and responds to a bout of exercise depending on its intensity (18,19). In this study, plasma corticosterone levels were measured 6 and 24 h after exercise, at which time points represent the late and early light periods, respectively (Fig. 1A). Plasma corticosterone commonly shows a higher value in the late light to early dark period, that is, before awakening (25). Consistent with previous studies, the average plasma corticosterone levels were higher at 6 h than 24 h after exercise. Although plasma corticosterone levels did not differ among the groups at either time point (Fig. 6A), its levels 6 h after exercise were positively correlated with percent changes in NEPA in both the ME and VE groups (Fig. 6B, C). Daily corticosterone release is regulated by many chronobiological mechanisms, such as clock-related genes, neuronal activity in the suprachiasmatic nucleus, and hypocretin neurons in the paraventricular nucleus (47,48). Therefore, our findings indicate that an increase in corticosterone levels before awakening could regulate NEPA levels following exercise.

Furthermore, in the present study, exercise intensity of the VE group (25 m/min) was higher than LT in mice (36). Considering that compensatory reduction in NEPA is specifically caused by vigorous exercise (Fig. 2E), compensatory reduction in NEPA following exercise may be associated with increased blood lactate levels. It was recently reported that activation of lactate receptors, known as hydroxycarboxylic acid receptor 1, decreases spontaneous locomotor activity in mice (49). Therefore, it is worth investigating whether lactate signaling is associated with a reduction in NEPA following vigorous exercise. As described above, further studies on corticosterone, lactate, or other molecules are required to better understand the underlying mechanism of compensatory reduction in NEPA following exercise.

To date, there is growing controversy over the effects of exercise programs on NEPA in human studies (28,50). A biological marker predicting post-exercise reduction in NEPA and other physiological activities is required to accurately serve an optimal exercise program to individuals for weight management. As an initial step in this inquiry, our identification of a vigorous exercise-specific reduction in NEPA and body temperature presents a promising animal model serving as a foundation for probing potential biomarkers. Therefore, we should in future investigate neuroendocrine mechanisms underlying a reduction in NEPA and body temperature following exercise. NEPA and thermogenesis are recognized to be regulated by multiple neuromodulators, for example, corticotropin-releasing hormone, neuropeptide Y, leptin, agouti-related protein, orexin, and ghrelin (51), which may be a potential target for investigating neuroendocrine mechanisms underlying post-exercise compensatory responses. Future investigations should aim to uncover these mechanisms, leading to the identification of peripheral biomarkers associated with neuroendocrine changes. Moreover, if the critical molecular changes pivotal in post-exercise reduction in NEPA can be detected in blood and saliva, it holds the potential to be applied in predicting changes in NEPA. This investigation would serve the consideration of an optimal exercise program tailored to individual responses.

### Implications of reduced NEPA following exercise on health-related outcomes

To date, moderate to vigorous physical activity is widely recognized as an effective strategy for improving physiological and mental health. However, vigorous exercise often causes excessive stress-related physiological responses, potentially diminishing the benefits of exercise (21,22). Consequently, moderate exercise is recognized as an optimal exercise program for improving health-related outcomes, especially brain function. Considering our findings of vigorous exercise-specific reductions in NEPA and body temperature (Fig. 2E, 3E), we postulated that the diminished benefits of vigorous exercise probably resulted from a reduction in NEPA. Given that NEPA refers to almost all activities in daily life, excluding structured and purposeful exercise, decreased NEPA likely leads people to physical inactivity, even if they engage in exercise. Lower levels of NEPA are closely linked to cognitive decline and dementia (8–10). Altogether, when we investigate which intensity of exercise is beneficial for overall health, it might be necessary to not only exercise itself but also to consider changes in other physiological activities after exercise.

A growing body of evidence demonstrates that light-intensity exercise is sufficient to improve brain function (52). We speculate that if light-intensity exercise increases subsequent NEPA, post-exercise NEPA may contribute to the light exercise-induced improvement of brain function. The exercise volume is very low during light exercise, but it is possible that light exercise positively affects post-exercise physiological activities, potentially resulting in facilitating health outcomes. Further investigation of the influence of several intensities, volumes, and types of exercise on post-exercise physiological activities, including NEPA, is required. Additionally, investigating the implications of changes in NEPA on health-related outcomes, such as learning and memory functions, as well as depression- and anxiety-like behaviors, is crucial for a comprehensive understanding of optimal exercise strategy for mental health.

### Perspectives and limitations

In the present study, we examined the effect of a single bout of exercise on NEPA and body temperature and demonstrated a vigorous exercise-specific reduction in NEPA and body temperature following exercise (Fig. 2E, 3E). Next, we should investigate whether repeated vigorous exercise chronically decreased NEPA and body temperature, and led to further body weight gain in mice. A previous study reported that a long-term extreme endurance event leads to a reduction in the energy expenditure generated by NEPA (53). This finding suggests that the vigorous exercise-induced reduction in NEPA continues over the long term. On the other hand, there is evidence that chronic exercise-induced enhancement of aerobic capacity is likely to increase NEPA (54). Therefore, comprehensive investigations focusing on exercise intensity, frequency, term, and changes in aerobic capacity are required to understand the optimal exercise programs for enhancing overall health.

In optimizing exercise interventions, careful consideration of the characteristics of individuals undergoing exercise programs is essential. It is pivotal to establish optimal exercise programs intended to both prevent obesity in non-obese individuals and improve obesity in those classified as obese. Given the importance of a negative energy balance in achieving body weight loss among obese individuals, specific attention would be required to be directed toward minimizing compensatory responses following exercise when prescribing exercise regimens. This underscores the need for an examination of the impact of exercise on NEPA and thermogenesis, with encompassing not only non-obese individuals but also those classified as obese. Therefore, in the subsequent phases of this study, it becomes critical to explore whether exercise induces a reduction in subsequent NEPA and thermogenesis in obese mice, with a consideration of understanding the influence of exercise intensity. This investigation would contribute to the refinement of exercise prescriptions tailored to the characteristics of individuals, developing effective strategies for both obesity prevention and management.

The effects of exercise are known to depend on the timing of exercise (55,56). In the current study, mice were subjected to exercise during the early light period, corresponding to the late afternoon or early night phase (Fig. 1A). Translating this timing to humans, it aligns with the period before sleep. In further investigation, it is imperative to scrutinize the effectiveness of different exercise timings, such as morning or noon, in preventing a reduction in NEPA and body temperature subsequent to exercise.

Our study demonstrated that plasma corticosterone levels were positively correlated with the percentage change in NEPA from pre-exercise to day 1 post-exercise (Fig. 6B, C). Although we suggested the possibility that insufficient pre-awakening corticosterone elevation contributes to a compensatory reduction in NEPA, it should be aware that corticosterone levels were analyzed using blood sampled after the exercise performed 3 days after the first exercise intervention as shown in the experimental procedure in Figure 1A. To prevent the sampling manipulation from confounding the effect of exercise on NEPA and body temperature, we performed the same exercise after measuring NEPA and body temperature and assessed corticosterone levels in response to exercise. Therefore, corticosterone levels after the first exercise are unknown. To better understand the association between daily corticosterone rhythms and compensatory reduction in NEPA following exercise, further examination using methods that enable noninvasive sampling of blood is required because our experimental design requires measurement of the physiological index after blood sampling.

## Conclusion

Our findings provide evidence for vigorous exercise-specific reductions in subsequent NEPA, body temperature, and their synchrony linked to weight gain. These compensatory responses to vigorous exercise must be induced by a disturbed circadian rhythm of corticosterone. Adjusting exercise intensity not only dictates the effectiveness of the exercise protocol itself, but also presents potential implications for post-exercise physical and/or physiological activities. This novel insight paves the way for a comprehensive exercise strategy in human life to address physical and mental health issues, mitigating the challenges of overweight and obesity beyond the energy expenditure of exercise itself.

## Author contributions

D.F. and T.M. conceived and designed the study. D.F., S.D., and K.S. performed the experiments. D.F. and T.M. analyzed the data. D.F., S.D., K.S., H.S., T.N., and T.M. interpreted the results of the experiments. D.F. and T.M. prepared the figures. D.F. and T.M. drafted the manuscript. All authors edited and revised the manuscript, and read and approved the final version of this manuscript.

## Data availability

The data that support the findings of this study are available from the corresponding author upon reasonable request.

## Acknowledgments

This study was supported by Grant-in-Aid for Scientific Research (B) (22H03478: Rep. T.M.), Grant-in-Aid for Scientific Research (C) (22K11528: Rep. T.N.), Grant-in-Aid for Research Activity Start-up (22K21199: Rep. D.F.) by JSPS KAKENHI, and by Japan Science and Technology Agency (JST) (JPMJFR205M: Rep. T.M.). The authors declare no competing interests. The results of the study are presented clearly, honestly, and without fabrication, falsification, or inappropriate data manipulation. The results of the present study do not constitute endorsement by the American College of Sports Medicine.

## Notes

### Competing Interest Statement

The authors have declared no competing interest.

### Summary of Updates

Title and main text updated to clarify our findings; Figure 4 revised.

## References

1. Levine JA, Lanningham-Foster LM, McCrady SK, et al. Interindividual variation in posture allocation: possible role in human obesity. Science. 2005;307(5709):584–6.

2. Stillman CM, Esteban-Cornejo I, Brown B, Bender CM, Erickson KI. Effects of exercise on brain and cognition across age groups and health states. Trends Neurosci. 2020;43(7):533–543.

3. Taylor JA, Greenhaff PL, Bartlett DB, Jackson TA, Duggal NA, Lord JM. Multisystem physiological perspective of human frailty and its modulation by physical activity. Physiol Rev. 2023;103(2):1137–1191.

4. Broskey NT, Martin CK, Burton JH, Church TS, Ravussin E, Redman LM. Effect of aerobic exercise-induced weight loss on the components of daily energy expenditure. Med Sci Sports Exerc. 2021;53(10):2164–2172.

5. Melanson EL. The effect of exercise on non-exercise physical activity and sedentary behavior in adults. Obes Rev. 2017;18(Suppl. 1):40–49.

6. Mansfeldt JM, Magkos F. Compensatory responses to exercise training as barriers to weight loss: changes in energy intake and non-exercise physical activity. Curr Nutr Rep. 2023;12(2):327–337.

7. Villablanca PA, Alegria JR, Mookadam F, Holmes DR Jr, Wright RS, Levine JA. Nonexercise activity thermogenesis in obesity management. Mayo Clin Proc. 2015;90(4):509–19.

8. Abbott RD, White LR, Ross GW, Masaki KH, Curb JD, Petrovitch H. Walking and dementia in physically capable elderly men. JAMA. 2004;292(12):1447–53.

9. Barnes DE, Blackwell T, Stone KL, Goldman SE, Hillier T, Yaffe K. Cognition in older women: the importance of daytime movement. J Am Geriatr Soc. 2008;56(9):1658–64.

10. Krell-Roesch J, Syrjanen JA, Vassilaki M, et al. Association of non-exercise physical activity in mid- and late-life with cognitive trajectories and the impact of APOE ε4 genotype status: the Mayo Clinic Study of Aging. Eur J Ageing. 2019;16(4):491–502.

11. Riou MÈ, Jomphe-Tremblay S, Lamothe G, et al. Energy compensation following a supervised exercise intervention in women living with overweight/obesity is accompanied by an early and sustained decrease in non-structured physical activity. Front Physiol. 2019;10:1048.

12. Drenowatz C, Grieve GL, DeMello MM. Change in energy expenditure and physical activity in response to aerobic and resistance exercise programs. Springerplus. 2015;4(1):798.

13. Liu XM, Wang K, Zhu Z, Cao ZB. Compensatory effects of different exercise durations on non-exercise physical activity, appetite, and energy intake in normal weight and overweight adults. Front Physiol. 2022;13:932846.

14. de Carvalho FP, Benfato ID, Moretto TL, Barthichoto M, de Oliveira CA. Voluntary running decreases nonexercise activity in lean and diet-induced obese mice. Physiol Behav. 2016;165:249–56.

15. Lark DS, Kwan JR, McClatchey PM, et al. Reduced nonexercise activity attenuates negative energy balance in mice engaged in voluntary exercise. Diabetes. 2018;67(5):831–840.

16. O’Neal TJ, Friend DM, Guo J, Hall KD, Kravitz AV. Increases in physical activity result in diminishing increments in daily energy expenditure in mice. Curr Biol. 2017;27(3):423–430.

17. Quintanilha ACS, Benfato ID, Santos RLO, Antunes HKM, de Oliveira CAM. Effects of acute exercise on spontaneous physical activity in mice at different ages. BMC Sports Sci Med Rehabil. 2021;13(1):78.

18. Morikawa R, Kubota N, Amemiya S, Nishijima T, Kita I. Interaction between intensity and duration of acute exercise on neuronal activity associated with depression-related behavior in rats. J Physiol Sci. 2021;71(1):1.

19. Nishii A, Amemiya S, Kubota N, Nishijima T, Kita I. Adaptive changes in the sensitivity of the dorsal raphe and hypothalamic paraventricular nuclei to acute exercise, and hippocampal neurogenesis may contribute to the antidepressant effect of regular treadmill running in rats. Front Behav Neurosci. 2017;11:235.

20. Jelleyman C, Yates T, O’Donovan G, et al. The effects of high-intensity interval training on glucose regulation and insulin resistance: a meta-analysis. Obes Rev. 2015;16(11):942–61.

21. Wu Y, Deng F, Wang J, et al. Intensity-dependent effects of consecutive treadmill exercise on spatial learning and memory through the p-CREB/BDNF/NMDAR signaling in hippocampus. Behav Brain Res. 2020;386:112599.

22. Shahroodi A, Mohammadi F, Vafaei AA, Miladi-Gorji H, Bandegi AR, Rashidy-Pour A. Impact of different intensities of forced exercise on deficits of spatial and aversive memory, anxiety-like behavior, and hippocampal BDNF during morphine abstinence period in male rats. Metab Brain Dis. 2020;35(1):135–147.

23. Otsuka T, Nishii A, Amemiya S, Kubota N, Nishijima T, Kita I. Effects of acute treadmill running at different intensities on activities of serotonin and corticotropin-releasing factor neurons, and anxiety- and depressive-like behaviors in rats. Behav Brain Res. 2016;298(Pt B):44–51.

24. Soya H, Nakamura T, Deocaris CC, et al. BDNF induction with mild exercise in the rat hippocampus. Biochem Biophys Res Commun. 2007;358(4):961–7.

25. Atkinson HC, Waddell BJ. Circadian variation in basal plasma corticosterone and adrenocorticotropin in the rat: sexual dimorphism and changes across the estrous cycle. Endocrinology. 1997;138(9):3842–8.

26. Dalm S, Enthoven L, Meijer OC, et al. Age-related changes in hypothalamic-pituitary-adrenal axis activity of male C57BL/6J mice. Neuroendocrinology. 2005;81(6):372–80.

27. Girard I, Garland T Jr. Plasma corticosterone response to acute and chronic voluntary exercise in female house mice. J Appl Physiol (1985). 2002; 92(4):1553–61.

28. Drenowatz C. Reciprocal compensation to changes in dietary intake and energy expenditure within the concept of energy balance. Adv Nutr. 2015;6(5):592–9.

29. Monnard CR, Fares EJ, Calonne J, et al. Issues in continuous 24-h core body temperature monitoring in humans using an ingestible capsule telemetric sensor. Front Endocrinol (Lausanne). 2017;8:130.

30. Manthou E, Gill JM, Wright A, Malkova D. Behavioral compensatory adjustments to exercise training in overweight women. Med Sci Sports Exerc. 2010;42(6):1121–8.

31. Thomas JV, Tobin SY, Mifflin MG, et al. The Effects of an Acute Bout of Aerobic or Resistance Exercise on Nonexercise Physical Activity. Exerc Sport Mov. 2023;1(2):e00004.

32. Reid KJ, Kräuchi K, Grimaldi D, et al. Effects of manipulating body temperature on sleep in postmenopausal women. Sleep Med. 2021;81:109–115.

33. Yamasue K, Hagiwara H, Tochukibo O, Sugimoto C, Kohno R. Measurement of core body temperature by an ingestible capsule sensor and evaluation of its wireless communication performance. Adv Biomed Eng. 2012;1:9–15.

34. Funabashi D, Wakiyama Y, Muto N, Kita I, Nishijima T. Social isolation is a direct determinant of decreased home-cage activity in mice: A within-subjects study using a body-implantable actimeter. Exp Physiol. 2022;107(2):133–146.

35. Kaneda Y, Kawata A, Suzuki K, Matsunaga D, Yasumatsu M, Ishiwata T. Comparison of neurotransmitter levels, physiological conditions, and emotional behavior between isolation-housed rats with group-housed rats. Dev Psychobiol. 2021;63(3):452–460.

36. Yook JS, Rakwal R, Shibato J, et al. Leptin in hippocampus mediates benefits of mild exercise by an antioxidant on neurogenesis and memory. Proc Natl Acad Sci U S A. 2019;116(22):10988–10993.

37. Pontzer H, Durazo-Arvizu R, Dugas LR, et al. Constrained total energy expenditure and metabolic adaptation to physical activity in adult humans. Curr Biol. 2016;26(3):410–7.

38. Abreu-Vieira G, Xiao C, Gavrilova O, Reitman ML. Integration of body temperature into the analysis of energy expenditure in the mouse. Mol Metab, 2015;4(6):461–70.

39. Zitting KM, Vujovic N, Yuan RK, et al. Human resting energy expenditure varies with circadian phase. Curr Biol. 2018;28(22):3685–3690.e3.

40. Škop V, Guo J, Liu N, et al. The metabolic cost of physical activity in mice using a physiology-based model of energy expenditure. Mol Metab. 2023;71:101699.

41. Nakamura TJ, Nakamura W, Yamazaki S, et al. Age-related decline in circadian output. J Neurosci. 2011;31(28):10201–5.

42. Yi Lee PM, Ling Kwok BH, Ting Ma JY, Tse LA. A population-based prospective study on rest-activity rhythm and mild cognitive impairment among Hong Kong healthy community-dwelling older adults. Neurobiol Sleep Circadian Rhythms. 2021;10:100065.

43. Yoon IY, Kripke DF, Elliott JA, Youngstedt SD, Rex KM, Hauger RL. Age-related changes of circadian rhythms and sleep-wake cycles. J Am Geriatr Soc. 2003;51(8):1085–91.

44. Hood S, Amir S. The aging clock: circadian rhythms and later life. J Clin Invest. 2017;127(2):437–446.

45. Oishi K, Yamamoto S, Uchida D, Doi R. Ketogenic diet and fasting induce the expression of cold-inducible RNA-binding protein with time-dependent hypothermia in the mouse liver. FEBS Open Bio. 2013;3:192–5.

46. Yasumoto Y, Nakao R, Oishi K. Free access to a running-wheel advances the phase of behavioral and physiological circadian rhythms and peripheral molecular clocks in mice. PLoS One. 2015;10(1):e0116476.

47. Bonnavion P, Jackson AC, Carter ME, de Lecea L. Antagonistic interplay between hypocretin and leptin in the lateral hypothalamus regulates stress responses. Nat Commun. 2015;6:6266.

48. Jones JR, Chaturvedi S, Granados-Fuentes D, Herzog ED. Circadian neurons in the paraventricular nucleus entrain and sustain daily rhythms in glucocorticoids. Nat Commun. 2021;12(1):5763.

49. Li J, Xia Y, Xu H, et al. Activation of brain lactate receptor GPR81 aggravates exercise-induced central fatigue. Am J Physiol Regul Integr Comp Physiol. 2022;323(5):R822–R831.

50. Silva AM, Júdice PB, Carraça EV, King N, Teixeira PJ, Sardinha LB. What is the effect of diet and/or exercise interventions on behavioral compensation in non-exercise physical activity and related energy expenditure of free-living adults? A systematic review. Br J Nutr. 2018;119(12):1327–1345.

51. Teske JA, Billington CJ, Kotz CM. Neuropeptidergic mediators of spontaneous physical activity and non-exercise activity thermogenesis. Neuroendocrinology. 2008;87(2):71–90.

52. Jesmin S, Shima T, Soya M, et al. Long-term light and moderate exercise intervention similarly prevent both hippocampal and glycemic dysfunction in presymptomatic type 2 diabetic rats. Am J Physiol Endocrinol Metab. 2022;322(3):E219–E230.

53. Thurber C, Dugas LR, Ocobock C, Carlson B, Speakman JR, Pontzer H. Extreme events reveal an alimentary limit on sustained maximal human energy expenditure. Sci Adv. 2019;5(6):eaaw0341.

54. Chow LS, Greenlund LJ, Asmann YW, et al. Impact of endurance training on murine spontaneous activity, muscle mitochondrial DNA abundance, gene transcripts, and function. J Appl Physiol (1985). 2007;102(3):1078–89.

55. Sasaki H, Miyakawa H, Watanabe A, et al. Evening rather than morning increased physical activity alters the microbiota in mice and is associated with increased body temperature and sympathetic nervous system activation. Biochim Biophys Acta Mol Basis Dis. 2022;1868(6):166373.

56. Sato S, Basse AL, Schönke M, et al. Time of exercise specifies the impact on muscle metabolic pathways and systemic energy homeostasis. Cell Metab. 2019;30(1):92–110.e4.

